# Spatially clustered resources increase aggregation and mating duration in *Drosophila melanogaster*

**DOI:** 10.1101/2020.05.06.080853

**Authors:** Emily R. Churchill, Jon R. Bridle, Michael D. F. Thom

## Abstract

In environments where females mate multiply, males should adjust their behaviour and physiology in response to the prevailing perceived level of sperm competition. This expectation is well supported by both laboratory and field studies, but we don’t yet know what mechanisms facilitate these plastic responses in natural populations. One way in which males appear to assess sperm competition risk is through encounter rates with conspecific males. Such encounter rates may be driven by the spatial distribution of resources required by male. However, explicit links between resource distribution, male encounter rate, and shifts in behaviour related to sperm competition have not been demonstrated. Here we show that a small increase in the distance of patches of food resources in the laboratory: (a) approximately halves the mean distances between pairs of males; and (b) is associated with an increase in subsequent copulation duration – previously shown to be a reliable indicator of male perception of sperm competition risk – by more than two minutes. Aggregation of resources, operating via increased encounter rate, is a mechanism that can stimulate plastic male sperm competition responses. Because spatial distribution of resources, including those exploited by *Drosophila*, is variable in nature, this may explain one way in which sperm competition-related plasticity is influenced in wild-living males.

## Introduction

Variation in population density affects the rate at which individuals encounter conspecific competitors and potential mates, with knock on consequences for the strength of sexual selection. One source of variation in local population density is the spatial structure of critical resources – clumped resources lead to increased encounter rates with competitors and mates as they gather to access those resources (Emlen and Oring, 1977). One adaptive response to encounter rate that has received considerable attention is the effect on investment in pre- and post-copulatory processes: with increasing encounter rate, these should be upregulated to maximise reproductive success in the new social environment (Kokko and Rankin, 2006). Several empirical studies have supported this prediction, including in crickets (Gage and Barnard, 1996), beetles (McCullough et al., 2018), bugs (García-González and Gomendio, 2004), platyhelminths (Giannakara et al., 2016), fish (Candolin and Reynolds, 2002), and rodents (Firman et al., 2018; Ramm and Stockley, 2009).

Demonstrations that male encounter rate can stimulate plasticity in sexual traits has generally been achieved by housing males at varying densities in the laboratory, with the most common treatment comparing a singly-housed male with a male housed with one or more conspecifics (Candolin and Reynolds, 2002; Firman et al., 2018; Gage and Barnard, 1996; Lizé et al., 2012; Moatt et al., 2013). This extreme manipulation of the total number of potential rivals is not intended to mimic the effects males experience in nature, but rather to demonstrate that such adaptive responses exist. Evidence for how such responses link to more ecologically-realistic stimuli is lacking, although effects of sperm competition have been observed in natural populations – for example in lizards (Kustra et al., 2019) and frogs (Buzatto et al., 2015). Given that patchiness in food resources is common in nature, and that resource distribution affects the degree of male-male competition (Emlen and Oring, 1977), small-scale variation in resource distribution that leads to local variation in encounter rate could drive the plastic effects in allocation of resources to sexual behaviour described above.

Laboratory studies have repeatedly demonstrated that *Drosophila melanogaster (Drosophilidae Diptera)* males are highly sensitive to the presence of other males, and that they increase their investment in sperm quality and ejaculate size (Garbaczewska et al., 2013; Hopkins et al., 2019; Moatt et al., 2014), investment in ejaculate composition (Fedorka et al., 2011; Hopkins et al., 2019; Wigby et al., 2009), and lengthen copulation durations (Bretman et al., 2009) when they perceive an elevated risk of sperm competition. Because *D. melanogaster* feed and breed on fermenting fruit (Begon, 1982), they rely on an inherently patchy resource with individual fruits naturally varying in size and proximity. Sex ratio and local population density of natural populations can vary considerably as a result (Markow, 1988; Soto-Yéber et al., 2018). This patchiness in natural food resources seems an ideal candidate for the type of ecological variability that might stimulate adjustment in post-copulatory processes in the wild.

West whether sperm competition-linked responses respond to resource patchiness by exposing male *D. melanogaster* to three different food distributions (clustered, dispersed and an even coverage control). In this way we can manipulate local density in an ecologically-realistic way, but without manipulating the number of rivals as previous laboratory studies have done (Bretman et al., 2009; Fedorka et al., 2011; Garbaczewska et al., 2013; Hopkins et al., 2019; Moatt et al., 2014; Wigby et al., 2009). We use the duration of copulation as a proxy for males’ perception of sperm competition risk, an association that has been demonstrated repeatedly in the laboratory (Bretman et al., 2009; Bretman et al., 2010; Bretman et al., 2012; Bretman et al., 2013; Mazzi et al., 2009; Moatt et al., 2013). We predict that: (a) by experimentally manipulating the distribution of food resources, males on clustered resources have a higher mean proximity to rivals (i.e. higher encounter rate), and (b) males on patchy resources will subsequently mate for longer indicating a perception of increased sperm competition risk.

## Methods

All fly rearing and experiments were conducted in a 12hour light:dark cycle (08:00 – 20:00h GMT), at 25°C. *Drosophila melanogaster* used were from a laboratory population (Canton-S), and populations were cultured on 7ml of a standard agar-based medium of 40g of yeast per litre, in 40ml vials. Between 20 and 30 *Drosophila* were housed in each vial. To minimise any effects of inbreeding, drift, and selective sweeps, every seven days the adults from all vials were pooled and randomly redistributed among new vials to start the next generation.

Test flies (180 in total – 60 per treatment) were collected from parent vials, each established with six males and six females allowed to breed for 70-98h. Test flies were removed from parent vials within six hours of eclosion to ensure virginity, and immediately aspirated under light ice anaesthesia into treatments. Virgin female flies for mating assays were collected from the same parental vials and aspirated into new vials in groups of four. Females were used in mating assays when they were seven days (+ 6-8 hours) old.

### Manipulating resource distributions and patchiness

Each replicate for each treatment consisted of four virgin males maintained in a 90mm Petri dish for three days. Food in each of these 45 dishes was arranged in one of three treatments (N = 15): clustered, dispersed or uniform (control) food resource distributions. Clustered and dispersed treatments both contained four plugs (420mm^3^ per patch) of standard food medium (as described above). The size of these patches is within the range of patch sizes where territorial behaviours have previously been observed (Hoffmann and Cacoyianni, 1990).

Dispersed food discs were placed at four equidistant points around the circumference of the Petri dish; these were 50mm apart along the edge of the square, 70mm apart on the diagonal (illustrated in Fig. 2). Clustered discs were placed in the centre of the Petri dish, in a square arrangement with each food disc in direct contact with adjacent discs. The control treatment was an even layer of 45ml standard medium covering the bottom of the dish (to the same height as the four food patches in the previous two treatments): volume and surface area were both greater in the control than the two clustered treatments, but given the number of flies food was assumed to be available *ad libitum* in all. All treatments were maintained in 12L:12D at 25°C, and the four male flies per treatment remained in these conditions for 70 hours (+/− 1hour) until aged to three days.

### Quantifying male spacing behaviour

Treatment enclosures were placed in one of two identical incubators maintained at 25°C and on the same 12:12 L:D cycle as the stock flies. Each incubator was fitted with a Raspberry Pi (www.raspberrypi.org) connected to an 8MP Raspberry Pi Camera module (v2; www.thepihut.com). Two to three Petri dishes, placed in a balanced arrangement across all treatment combinations, were placed directly under each camera. We used frame capture software (‘raspistill’) to collect one image every 15 minutes from 08:00-20:00 GMT (during the light part of the cycle). We captured the x-y coordinates of each male at each time point using ImageJ’s multiple point selector tool (Schneider et al., 2012), and then converted these into a set of six Euclidean pairwise distances between the four males (24670 measurements across the three treatments and all time points). For 325 out of the 4290 individual time-point photographs (7.6%) we were unable to accurately locate at least one male on the image. To minimize the effect of missing data on the number of time points included per replicate, the unit of analysis was the mean (rather than the raw data) of the distances between each pair for each time point.

### Reproductive behavioural assays

After 70h in treatment, each male from each Petri dish was allowed one opportunity to mate with a standard seven day old (±3h) virgin female and mating behaviours were observed (N = 15; 60 individuals). The male and female were aspirated into a standard food vial supplemented with ~0.03g active yeast granules. The space in the vial was limited to 7cm3 by pushing the vial bung down into the vial to reduce encounter latency. Courtship latency was defined as the time from which the pair were first introduced until the male initiated his first wing extension. Latency to copulate (courtship duration) started at the time of the first wing extension, and ended with a male’s successful mounting attempt. Copulation duration was recorded from successful mounting until the pair were fully separated.

Not every male courted (control: 81.8%; clustered: 86.4%; dispersed: 95.6%), and not all courting males mated (control: 75.0%; clustered: 86.8%; dispersed: 83.3%). We observed each pair for a maximum of 90 minutes after the pair had been introduced, and recorded failure to court and/or failure to mate after this time.

### Statistical analysis

Sample sizes were 15 replicates (N = 60 *Drosophila*) for each of the three treatments, of which 11 from each treatment (33 in total) were photographed to collect spacing data. The effect of treatment on total inter-male distance was analysed using linear mixed effects models, with plate and time point modelled as random effects.

Treatment effects on mating related traits were analysed using linear mixed effects models, with replicate plate (nested within treatment) entered as a random effect to account for the fact that mating data was available for (up to) four males per plate. Time point (numbered sequentially from first to last measurement and treated as continuous) and treatment were initially entered as interacting predictor variables; if the interaction was non-significant we re-ran the model with both variables entered as main effects. We used the R package lmerTest (Kuznetsova et al., 2017) to generate p values using the Satterthwaite approximation for degrees of freedom. To assess the effect of treatment on binomial variables (courtship success, copulation success) we used generalised linear mixed models with a binomial error distribution, and replicate plate nested within treatment to account for possible plate effects.

## Results

### Effect of food distribution on inter-male spacing

The spatial distribution of food patches significantly affected the mean pairwise distance between the four individuals in the population (treatment*time: F_2,4239_ = 286, p << 0.001; Fig. 1). On the final day of exposure to food treatment, pairwise distance between males in the dispersed treatment (44.02 ± 0.66mm) was nearly twice the pairwise distance in the clustered food treatment (22.79 ± 0.86mm; main effect of treatment: F_1,20_ = 57.8, p << 0.001). Control males were also significantly further apart from one another on the final day (mean pairwise distance 39.35±0.93) than pairs of flies from the clustered treatment (F_1,20_ = 27.9, p < 0.001), but they were not significantly different in mean inter-male distance than those in the dispersed treatment (F_1,20_ = 3.9, p = 0.061).

**Figure 1.**
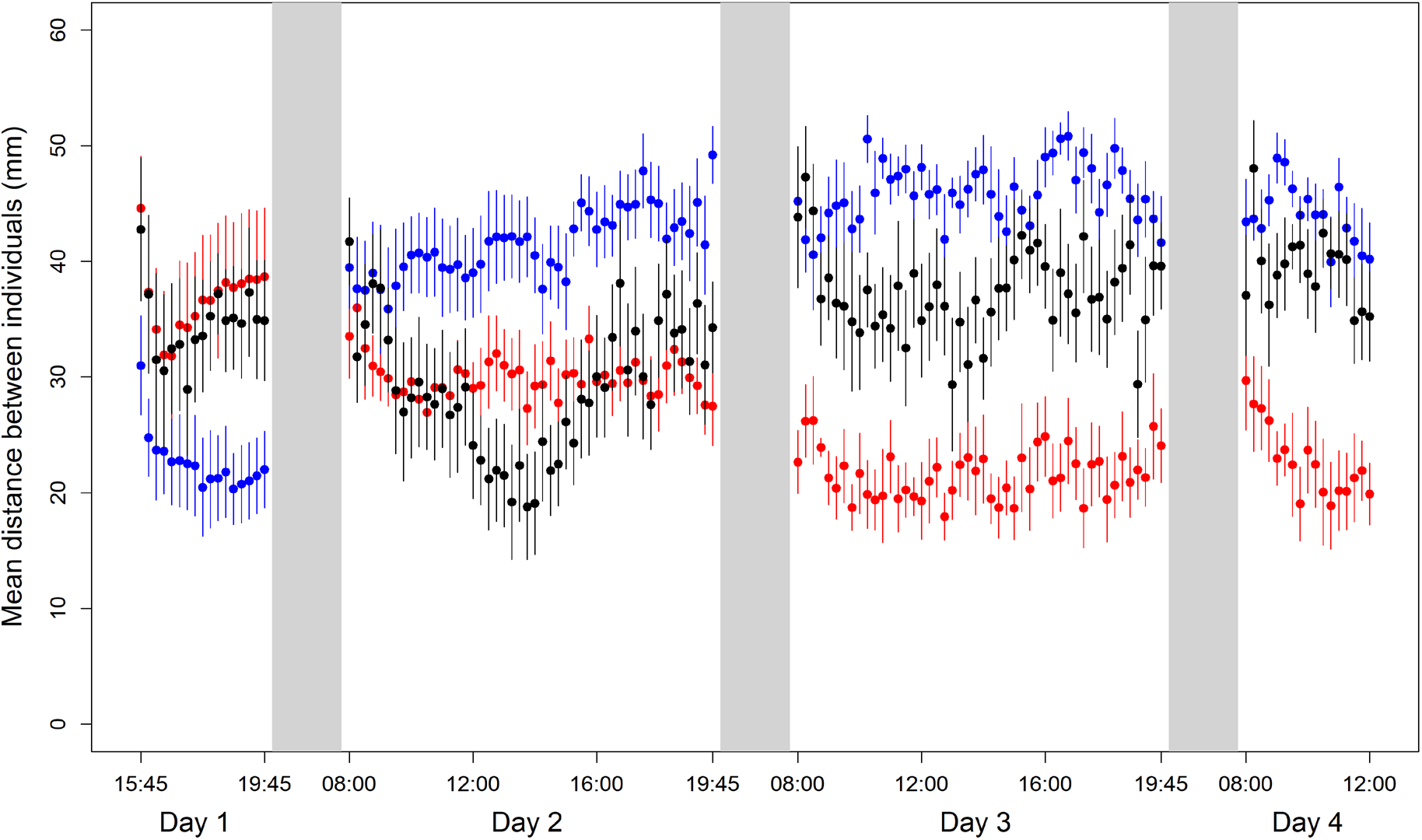
Mean inter-fly distance (mean of 6 pairwise distances between 4 focal flies per plate, averaged across 11 replicate plates) over time. Black = control treatment (evenly distributed food); red = clustered food patches; blue = dispersed food patches. Bars show standard errors of the mean for each time point across all 11 treatment replicates. Grey blocks indicate period of dark (20:00 - 08:00h GMT), and are not to scale.

### Effect of resource distribution on mating behaviour

Among those males that mated, copulation duration was significantly affected by food distribution (mixed effects model: F_2,40.9_ = 3.96, p = 0.026; Fig. 2). We re-tested the effect of treatment using the mean mating duration of all males from a replicate as the unit of analysis, finding the same result (F_2,42_ = 4.22, p = 0.021). Males from the clustered treatment mated for significantly longer (1170 ± 28s) than those from the dispersed treatment (1029 ± 28s), a difference of 2 minutes 20 seconds (mixed effects model: F_1,28_ = 6.59, p = 0.016). Copulation duration of males from the control treatment did not significantly differ from either of the other treatments (control copulation duration 1107 ± 23s; vs. dispersed: F_1,28.5_ = 2.22, p = 0.146; vs. clustered F_1,28.5_ = 1.96, p = 0.172). However, despite these observed differences between clustered and dispersed treatments, the mean distance between males did not significantly affect copulation duration in any of the three treatments (all p > 0.101).

**Figure 2.**
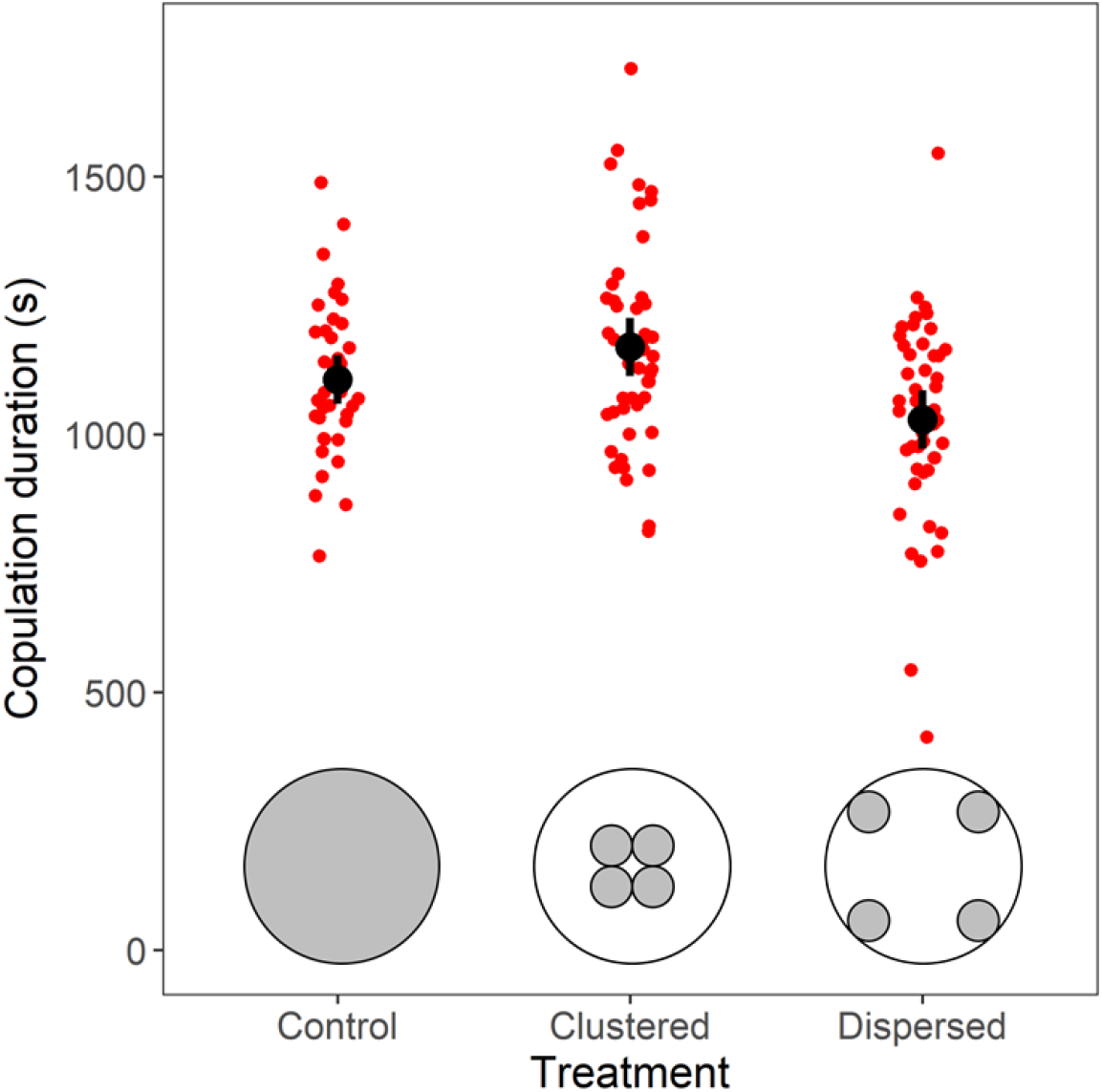
The effect of food resource spatial distribution on the duration of subsequent copulation. Means (black dot) and 95% confidence intervals of copulation duration (seconds). Sample sizes: clustered 49 (11 males did not mate), control 44 (16), dispersed 51 (9).

In total, 159 of 180 males (88.3%) courted the female. There was no significant effect of treatment on the proportion of males that courted (generalized linear model with binomial errors and plate nested within treatment; χ^2^ = 118, p = 0.376). Similarly, 144 (80%) of males mated, and this was not influenced by treatment (χ^2^ = 175, p = 0.286). Neither the latency to start courting (F_2,39.3_ = 0.201 p = 0.818) nor the latency to start copulation (F_2,30.4_ = 1.257, p = 0.299), differed significantly among the three treatments.

## Discussion

The high degree of plasticity in mating-related traits shown by male *Drosophila* is now well established (Churchill et al., 2019; Davies et al., 2019; Droney, 1998; Fricke et al., 2008; Jensen et al., 2015; Lefranc and Bungaard, 2000; Lüpold et al., 2010; Morimoto and Wigby, 2016; Ormerod et al., 2017; Schultzhaus et al., 2017). Variation in these traits is highly sensitive to conspecific male density in a manner which suggests that males adjust investment in anticipation of the intensity of sperm competition they are likely to encounter during mating (Bretman et al., 2009). However, how this level of plasticity relates to variation in density observed in natural populations remains unknown, and laboratory studies tend to manipulate density in ways that seem unlikely to occur frequently in nature (e.g. singly-housed males compared to a high density of males in a single vial).

We show that manipulating food patchiness while keeping group size constant has the same effect on a sperm competition-related trait – both in direction and magnitude – as manipulating local density directly, and that these effects can be observed even over very small spatial scales. As wild *D. melanogaster* encounter a patchy resource that is likely to alter male encounter rates at a similar scale to that demonstrated here (Markow, 1988; Soto-Yéber et al., 2018), we suggest that this is a mechanism by which the environment might influence male allocation of resources to traits associated with sperm competition, and thus mating success, in wild-living *Drosophila*.

As in previous studies, male *Drosophila* responded to an increased perceived sperm competition with a lengthened copulation duration (by over two minutes) when introduced to a mating partner (Bretman et al., 2009; Bretman et al., 2012). While the effect on mating duration is a repeatable indicator of male perception of sperm competition risk, the benefits of this behaviour to males remains unresolved. In many species, increased mating duration has been linked to increased sperm transfer and offspring production (Edvardsson and Canal, 2006; Engqvist and Sauer, 2003; Sakaluk and Eggert, 1996). In *Drosophila* the consequences of longer copulation durations are less clear, with some studies reporting an association with increased fitness (Bretman et al., 2009; Garbaczewska et al., 2013; Price et al., 2012), while others have not found a link (Bretman et al., 2012; Dobler and Reinhardt, 2016). Whether males on the clustered food resource would have a higher fitness than those on dispersed resources remains to be tested, but will almost certainly depend on mating order effects and the competing male’s history of exposure to rivals (Bretman et al., 2012). However, our objective here was not to examine fitness consequences, but rather to demonstrate that males apparently perceive effects on sperm competition risk that result directly from small-scale changes in the spatial distribution of resources.

Interestingly, the effect of food distribution on male behaviour was observed in the absence of females. Females often follow social cues, and their grouping behaviour is promoted by aggregation pheromones (Bartelt et al., 1985; Duménil et al., 2016).. By comparison, given their low feeding rate once adult (Wong et al., 2009), males are thought to aggregate near food resources primarily to seek mating opportunities. That males were responsive to the distribution of food even in the absence of females is intriguing, and the relative importance of female social cues and the direct response to food resources themselves are yet to be determined. In general, studies manipulating male density have tended to exclude females from the treatment phase (e.g. Bretman et al. (2009); Bretman et al. (2010); Lizé et al. (2012); Moatt et al. (2013); Price et al. (2012); and Rouse and Bretman (2016)), and the effects of inter-sexual interactions on plastic responses to density remains a relatively unexplored area.

This study adds to a small number of studies demonstrating the effect that environmental heterogeneity can have on *Drosophila* behaviour. Yun et al. (Yun et al., 2017) demonstrated that female fitness was higher in more spatially complex laboratory environments as a result of a reduction in sexual interactions and consequent mitigation of male harm. Similar effects had previously been demonstrated when laboratory populations were presented with a refuge: female remating rates declined substantially (Byrne et al., 2008). Such rapid shifts in behaviour, driven by ecological patchiness, have to date rarely been included in laboratory assays, but may have major effects on the demography and growth rate of populations exposed to spatial patchiness, through their effects on male reproductive skew and therefore effective population size. These effects may have important evolutionary and ecological consequences in relatively patchy parts of a species’ distribution, for example by increasing sexual conflict over shared resources (Pilakouta et al., 2016), or reducing maximum sustainable rates of evolution (Bridle et al., 2009).

There are some intriguing dynamics operating in the inter-male distances in the early stages of the treatment period: in particular, males on the dispersed food patches initially experience lower inter-male distances than those on the clustered food (Figure 1). This effect is does not match what we expected to see among males attempting to defend individual patches, and is the opposite to the pattern observed on the final days of treatment. Inspection of photographs from this treatment suggests that males on the dispersed food patches initially cluster together away from food before sorting themselves into individual territories focussed around each patch. Territorial behaviour in *D. melanogaster* has previously been observed under laboratory conditions, and appears to be driven by boundaries of food sources (Lim et al., 2014) so it is possible that multiple distinct territories could be established under these conditions. However, as yet it is not clear what is driving the initial clustering behaviour.

Our results demonstrate a clear link between small-scale patchiness of resources and behaviours that suggest male sensitivity to sperm competition risk, mediated by changes in male-male encounter rate. While density effects on male mating duration have been demonstrated several times, we have placed this response in a biologically meaningful context by demonstrating a link to ecological factors that are very likely to be at play in wild-living populations.

## References

Bartelt RJ, Schaner AM, Jackson LL, 1985. cis-Vaccenyl acetate as an aggregation pheromone in *Drosophila melanogaster*. Journal of Chemical Ecology 11:1747–1756. doi: 10.1007/BF01012124.

Begon M, 1982. Yeasts and *Drosophila*. The genetics and biology of Drosophila 3:3345–3384.

Bretman A, Fricke C, Chapman T, 2009. Plastic responses of male *Drosophila melanogaster* to the level of sperm competition increase male reproductive fitness. Proceedings of the Royal Society B: Biological Sciences 276:1705–1711. doi: 10.1098/rspb.2008.1878.

Bretman A, Fricke C, Hetherington P, Stone R, Chapman T, 2010. Exposure to rivals and plastic responses to sperm competition in *Drosophila melanogaster*. Behavioral Ecology 21:317–321. doi: 10.1093/beheco/arp189.

Bretman A, Westmancoat James D, Gage Matthew JG, Chapman T, 2012. Individual plastic responses by males to rivals reveal mismatches between behaviour and fitness outcomes. Proceedings of the Royal Society B: Biological Sciences 279:2868–2876. doi: 10.1098/rspb.2012.0235.

Bretman A, Westmancoat JD, Chapman T, 2013. Male control of mating duration following exposure to rivals in fruitflies. Journal of Insect Physiology 59:824–827. doi: 10.1016/j.jinsphys.2013.05.011.

Bridle J, Polechová J, Vines T, 2009. Limits to adaptation and patterns of biodiversity. p. 77–101.

Buzatto BA, Roberts JD, Simmons LW, 2015. Sperm competition and the evolution of precopulatory weapons: Increasing male density promotes sperm competition and reduces selection on arm strength in a chorusing frog. Evolution 69:2613–2624. doi: 10.1111/evo.12766.

Byrne PG, Rice GR, Rice WR, 2008. Effect of a refuge from persistent male courtship in the *Drosophila* laboratory environment. Integrative and Comparative Biology 48:e1–e1. doi: 10.1093/icb/icn001.

Candolin U, Reynolds JD, 2002. Adjustments of ejaculation rates in response to risk of sperm competition in a fish, the bitterling (*Rhodeus sericeus*). Proceedings of the Royal Society B: Biological Sciences 269:1549–1553. doi: 10.1098/rspb.2002.2055.

Churchill ER, Dytham C, Thom MDF, 2019. Differing effects of age and starvation on reproductive performance in *Drosophila melanogaster*. Scientific Reports 9:2167. doi: 10.1038/s41598-019-38843-w.

Davies LR, Schou MF, Kristensen TN, Loeschcke V, 2019. Fluctuations in nutrient composition affect male reproductive output in *Drosophila melanogaster*. Journal of Insect Physiology 118:103940. doi: 10.1016/j.jinsphys.2019.103940.

Dobler R, Reinhardt K, 2016. Heritability, evolvability, phenotypic plasticity and temporal variation in sperm-competition success of *Drosophila melanogaster*. Journal of Evolutionary Biology 29:929–941. doi: 10.1111/jeb.12858.

Droney DC, 1998. The influence of the nutritional content of the adult male diet on testis mass, body condition and courtship vigour in a Hawaiian *Drosophila*. Functional Ecology 12:920–928. doi: 10.1046/j.1365-2435.1998.00266.x

Duménil C, Woud D, Pinto F, Alkema JT, Jansen I, Van Der Geest AM, Roessingh S, Billeter J-C, 2016. Pheromonal cues deposited by mated females convey social information about egg-laying sites in *Drosophila melanogaster*. Journal of Chemical Ecology 42:259–269. doi: 10.1007/s10886-016-0681-3.

Edvardsson M, Canal D, 2006. The effects of copulation duration in the bruchid beetle *Callosobruchus maculatus*. Behavioral Ecology 17:430–434. doi: 10.1093/beheco/arj045.

Emlen ST, Oring LW, 1977. Ecology, sexual selection, and the evolution of mating systems. Science 197:215. doi: 10.1126/science.327542.

Engqvist L, Sauer KP, 2003. Determinants of sperm transfer in the scorpionfly *Panorpa cognata*: Male variation, female condition and copulation duration. Journal of Evolutionary Biology 16:1196–1204. doi: 10.1046/j.1420-9101.2003.00613.x.

Fedorka KM, Winterhalter WE, Ware B, 2011. Perceived sperm competition intensity influences seminal fluid protein production prior to courtship and mating. Evolution 65:584–590. doi: 10.1111/j.1558-5646.2010.01141.x.

Firman RC, Garcia-Gonzalez F, Simmons LW, André GI, 2018. A competitive environment influences sperm production, but not testes tissue composition, in house mice. Journal of Evolutionary Biology 31:1647–1654. doi: 10.1111/jeb.13360.

Fricke C, Bretman A, Chapman T, 2008. Adult male nutrition and reproductive success in *Drosophila melanogaster*. Evolution 62:3170–3177. doi: 10.1111/j.1558-5646.2008.00515.x.

Gage AR, Barnard CJ, 1996. Male crickets increase sperm number in relation to competition and female size. Behavioral Ecology and Sociobiology 38:349–353. doi: 10.1007/s002650050251.

Garbaczewska M, Billeter J-C, Levine JD, 2013. *Drosophila melanogaster* males increase the number of sperm in their ejaculate when perceiving rival males. Journal of Insect Physiology 59:306–310. doi: 10.1016/j.jinsphys.2012.08.016.

García-González F, Gomendio M, 2004. Adjustment of copula duration and ejaculate size according to the risk of sperm competition in the golden egg bug (*Phyllomorpha laciniata*). Behavioral Ecology 15:23–30. doi: 10.1093/beheco/arg095.

Giannakara A, Schärer L, Ramm SA, 2016. Sperm competition-induced plasticity in the speed of spermatogenesis. BMC Evolutionary Biology 16:60. doi: 10.1186/s12862-016-0629-9.

Hoffmann AA, Cacoyianni Z, 1990. Territoriality in *Drosophila melanogaster* as a conditional strategy. Animal Behaviour 40:526–537. doi: 10.1016/S0003-3472(05)80533-0.

Hopkins BR, Sepil I, Thézénas M-L, Craig JF, Miller T, Charles PD, Fischer R, Kessler BM, Bretman A, Pizzari T, Wigby S, 2019. Divergent allocation of sperm and the seminal proteome along a competition gradient in *Drosophila melanogaster*. Proceedings of the National Academy of Sciences:201906149. doi: 10.1073/pnas.1906149116.

Jensen K, McClure C, Priest NK, Hunt J, 2015. Sex-specific effects of protein and carbohydrate intake on reproduction but not lifespan in *Drosophila melanogaster*. Aging cell 14:605–615. doi: 10.1111/acel.12333.

Kokko H, Rankin DJ, 2006. Lonely hearts or sex in the city? Density-dependent effects in mating systems. Philosophical Transactions of the Royal Society B: Biological Sciences 361:319–334. doi: 10.1098/rstb.2005.1784.

Kustra MC, Kahrl AF, Reedy AM, Warner DA, Cox RM, 2019. Sperm morphology and count vary with fine-scale changes in local density in a wild lizard population. Oecologia 191:555–564. doi: 10.1007/s00442-019-04511-z.

Kuznetsova A, Brockhoff PB, Christensen RHB, 2017. lmerTest Package: Tests in Linear Mixed Effects Models. Journal of Statistical Software 1. doi: 10.18637/jss.v082.i13.

Lefranc A, Bungaard J, 2000. The influence of male and female body size on copulation duration and fecundity in *Drosophila melanogaster*. Hereditas 132:243–247. doi: 10.1111/j.1601-5223.2000.00243.x.

Lim RS, Eyjólfsdóttir E, Shin E, Perona P, Anderson DJ, 2014. How food controls aggression in *Drosophila*. PLOS ONE 9:e105626. doi: 10.1371/journal.pone.0105626.

Lizé A, Price TAR, Marcello M, Smaller EA, Lewis Z, Hurst GDD, 2012. Males do not prolong copulation in response to competitor males in the polyandrous fly *Drosophila bifasciata*. Physiological Entomology 37:227–232. doi: 10.1111/j.1365-3032.2012.00836.x.

Lüpold S, Manier MK, Ala-Honkola O, Belote JM, Pitnick S, 2010. Male *Drosophila melanogaster* adjust ejaculate size based on female mating status, fecundity, and age. Behavioral Ecology 22:184–191. doi: 10.1093/beheco/arq193.

Markow TA, 1988. Reproductive behavior of *Drosophila melanogaster* and *D. nigrospiracula* in the field and in the laboratory. Journal of Comparative Psychology 102:169–173. doi: 10.1037/0735-7036.102.2.169.

Mazzi D, Kesäniemi J, Hoikkala A, Klappert K, 2009. Sexual conflict over the duration of copulation in *Drosophila montana*: Why is longer better? BMC Evolutionary Biology 9:132. doi: 10.1186/1471-2148-9-132.

McCullough EL, Buzatto BA, Simmons LW, 2018. Population density mediates the interaction between pre- and postmating sexual selection. Evolution 72:893–905. doi: 10.1111/evo.13455.

Moatt JP, Dytham C, Thom MD, 2013. Exposure to sperm competition risk improves survival of virgin males. Biol Lett 9:20121188. doi: 10.1098/rsbl.2012.1188.

Moatt JP, Dytham C, Thom MD, 2014. Sperm production responds to perceived sperm competition risk in male *Drosophila melanogaster*. Physiol Behav 131:111–114. doi: 10.1016/j.physbeh.2014.04.027.

Morimoto J, Wigby S, 2016. Differential effects of male nutrient balance on pre- and post-copulatory traits, and consequences for female reproduction in *Drosophila melanogaster*. Sci Rep 6:27673. doi: 10.1038/srep27673.

Ormerod KG, LePine OK, Abbineni PS, Bridgeman JM, Coorssen JR, Mercier AJ, Tattersall GJ, 2017. *Drosophila* development, physiology, behavior, and lifespan are influenced by altered dietary composition. Fly 11:153–170. doi: 10.1080/19336934.2017.1304331.

Pilakouta N, Richardson J, Smiseth PT, 2016. If you eat, I eat: Resolution of sexual conflict over consumption from a shared resource. Animal Behaviour 111:175–180. doi: 10.1016/j.anbehav.2015.10.016.

Price TAR, Lizé A, Marcello M, Bretman A, 2012. Experience of mating rivals causes males to modulate sperm transfer in the fly *Drosophila pseudoobscura*. Journal of Insect Physiology 58:1669–1675. doi: 10.1016/j.jinsphys.2012.10.008.

Ramm SA, Stockley P, 2009. Adaptive plasticity of mammalian sperm production in response to social experience. Proceedings of the Royal Society B: Biological Sciences 276:745–751. doi: 10.1098/rspb.2008.1296.

Rouse J, Bretman A, 2016. Exposure time to rivals and sensory cues affect how quickly males respond to changes in sperm competition threat. Animal Behaviour 122:1–8. doi: 10.1016/j.anbehav.2016.09.011.

Sakaluk SK, Eggert A-K, 1996. Female control of sperm transfer and intraspecific variation in sperm precedence: Antecedents to the evolution of a courtship food gift. Evolution 50:694–703. doi: 10.1111/j.1558-5646.1996.tb03879.x.

Schneider CA, Rasband WS, Eliceiri KW, 2012. NIH Image to ImageJ: 25 years of image analysis. Nature Methods 9:671. doi: 10.1038/nmeth.2089.

Schultzhaus JN, Nixon JJ, Duran JA, Carney GE, 2017. Diet alters *Drosophila melanogaster* mate preference and attractiveness. Animal Behaviour 123:317–327. doi: 10.1016/j.anbehav.2016.11.012.

Soto-Yéber L, Soto-Ortiz J, Godoy P, Godoy-Herrera R, 2018. The behavior of adult *Drosophila* in the wild. PLOS ONE 13:e0209917. doi: 10.1371/journal.pone.0209917.

Wigby S, Sirot LK, Linklater JR, Buehner N, Calboli FCF, Bretman A, Wolfner MF, Chapman T, 2009. Seminal fluid protein allocation and male reproductive success. Current Biology 19:751–757. doi: 10.1016/j.cub.2009.03.036.

Wong R, Piper MDW, Wertheim B, Partridge L, 2009. Quantification of food intake in *Drosophila*. PLOS ONE 4:e6063. doi: 10.1371/journal.pone.0006063.

Yun L, Chen PJ, Singh A, Agrawal AF, Rundle HD, 2017. The physical environment mediates male harm and its effect on selection in females. Proc Biol Sci 284. doi: 10.1098/rspb.2017.0424.

